# 3D Functional Ultrasound Imaging of Pigeons

**DOI:** 10.1101/302323

**Authors:** Richard Rau, Pieter Kruizinga, Frits Mastik, Markus Belau, Nico de Jong, Johannes G. Bosch, Wolfgang Scheffer, Georg Maret

**Author notes:** http://cms.uni-konstanz.de/physik/maret/ (Richard Rau).

## Abstract

Recent advances in ultrasound Doppler imaging have allowed to visualize brain activity in small mammalian species such as rats and mice. In birds, this type of functional ultrasound imaging was impossible up to now because birds have physiological characteristics that are unfavorable for current functional ultrasound acquisition schemes. Here, we introduce a high-definition functional ultrasound acquisition method (HDfUS) acquiring 20,000 frames per second continuously. This enabled first successful functional studies on awake pigeons subjected to auditory and visual stimulation. We show that the improved spatiotemporal resolution and sensitivity of HDfUS allows to visualize and investigate the temporally resolved 3D neural activity evoked by a complex stimulation pattern, such as a moving light source. This illustrates the enormous potential of HDfUS imaging to become a new standard functional brain imaging method revealing unknown, stimulus related hemodynamics at excellent signal-to-noise ratio and spatiotemporal resolution.

**Highlights:**

- We describe a novel ultrafast functional ultrasound technique (HDfUS)
- HDfUS offers continuous recording with unmatched spatiotemporal resolution
- HDfUS allows to resolve complex 4D neurovascular responses in the brain
- First fUS study on non-mammalian species

## Introduction

Birds are classical and favorite animal models in a number of neurobiological disciplines. Together with songbirds, pigeons are the most studied species within the field of avian neuroimaging, leading to findings on language learning, vocalization (Doupe and Kuhl, 1999; Jarvis, 2004; Van Ruijssevelt et al., 2013; Voss et al., 2007) as well as interhemispheric communication (Jung-Beeman, 2005; Manns and Römling, 2012), learning and cognition (Browning et al., 2011; Güntürkün et al., 2000).

However, studying the avian brain *in vivo* is very cumbersome and often lacks one or more of the key characteristics for functional neuroimaging. For instance, most functional studies on birds are based on electrophysiological recordings (Freund et al., 2016; Ng et al., 2010), which allow very fast and sensitive measurements of local brain responses but are lacking either spatial resolution or field of view. The gold standard in functional imaging for humans, functional magnetic resonance imaging (fMRI), is challenging in birds due to the smaller cerebral structure, which require higher magnetic fields (Niranjan et al., 2016) and impair both signal-to-noise ratio (SNR) and temporal resolution (Logothetis, 2008). In addition, the high investment and maintenance cost of a MRI apparatus in general and of dedicated small animal MRI in particular, combined with space constrains and limited accessibility to supporting facilities, make MRI an inconvenient choice for small animal functional studies. Other methods investigating the avian brain *in vivo*, such as voltage sensitive dyes (Ng et al., 2010), are rare and are lacking either spatiotemporal resolution, penetration depth or good signal-to-noise ratio.

Recently, it has been shown that images of the blood volume inside the brain microvasculature can be obtained through high frame rate Doppler ultrasound imaging (also called μ-Doppler) (Bercoff et al., 2011) for small mammals such as rats (Macé et al., 2013; Tanter and Fink, 2014). The high spatiotemporal resolution of ∼ (100 μm)^2^ every few seconds allows to resolve and monitor local blood volume changes due to neural activation inside specific brain areas of rodents (Macé et al., 2011). This method has been named functional ultrasound (fUS) and has led to several interesting functional studies, investigating the somatosensory (Macé et al., 2011; Osmanski et al., 2014; Tiran et al., 2017; Urban et al., 2014, 2015), visual (Gesnik et al., 2017) and olfactory (Osmanski et al. 2014-2, Rungta et al., 2017) system of rats/mice along with discussions on functional connectivity (Osmanski et al., 2014; Urban et al., 2015), epilepsy (Macé et al., 2011; Sieu et al., 2015) and awake measurements (Sieu et al., 2015; Tiran et al., 2017; Urban et al., 2015). Very recently its applicability to brain microvasculature in human neonates was shown by revealing different sleep states and congenital cortical abnormalies (Demené et al., 2017).

So far, all these notable achievements with functional US have concentrated on the cerebral dynamics of rodents and could not be transferred to birds due to their different physiology (Gunkel and Lafortune, 2005), i.e. mainly their unfavorably low heart rate (see Supplementary Note 1 for a detailed discussion). In order to generalize functional ultrasound imaging to species with much wider range of physiological characteristics (in particular heart beat), we developed a variant of fUS with improved spatiotemporal resolution and filtering which we term High Definition fUS, HDfUS. The key improvement of HDfUS over fUS is ultrafast continuous realtime data acquisition and processing. As a proof of concept, we show that HDfUS not only gives access to the investigation of the avian neurobiology, but also enables to study temporally complex stimulation patters in the pigeon brain. The study first focuses on 2D and 3D HDfUS with results of responses in the auditory system of pigeons. In the second part we analyze the activity evoked by visual stimulation with a static and a moving light source.

## Materials and Methods

### High definition functional ultrasound imaging principle

In fUS, the cerebral blood volume (CBV) is measured by means of Power Doppler imaging (Shung et al., 1976). In Power Doppler imaging, the Doppler frequency shift of the ultrasound wave reflected from moving red blood cells is exploited. In any point of the image, the signal power of the Doppler-shifted ultrasound is proportional to the volume of moving blood. A locally increasing blood volume induces a higher signal of the Doppler shifted ultrasound power which is reflected from the larger amount of moving erythrocytes in the imaged microvasculature. Similar to the blood-oxygen-level-dependent (BOLD) signal in fMRI, the local CBV is modulated by the activation of the neurons surrounding the vascularity due to the neurovascular coupling (Macé et al., 2013) which in turn results from the enhanced metabolism of the activated neurons. Naturally, the CBV is highly dependent on the physiological characteristics of the species. Contrary to rats and mice, the comparably slow heartbeat of pigeons is unfavorable in terms of conventional fUS measurements, which are based on a Power Doppler imaging framerate of < 2 Hz. Using a state-of-the-art ultrasound research system and specialized image reconstruction algorithms (Kruizinga et al., 2012) powered by graphics processing unit (GPU) computing we realized the HDfUS acquisition scheme with a Power Doppler imaging frame rate up to several hundred Hz (corresponding to a data acquisition rate of ∼3.3 GB/s continuously). This enables robust filtering of the heartbeat influence (see Supplementary Note 1 for a detailed analysis) and is at present unmatched in the field (Gesnik et al., 2017; Sieu et al., 2015). Along with the development of HDfUS, it was of major importance to establish a setup for pigeons that allows awake recordings with a robust fixation and provides ultrasound penetration of the brain tissue. In the Methods and Supplementary Material section we outline a full protocol that allows establishing a working setup for awake measurements with the same animal over weeks.

### Animals and Anesthesia

All experiments were performed in accordance with the regulations of the German Animal Welfare Act (TierSchG) and have been approved by the regional committee of animal welfare (35-9185.81/G-16/99 and 35-9185.81/G-12/54). The HDfUS studies were carried out on three pigeons (columba livia domestica, one female, two male), one anesthetized and two awake. Anesthesia was performed with the pigeon breathing in a half open mask with pure oxygen and isoflurane supply. The isoflurane concentration is controlled by the Eickemeyer Anesthetic Unit Research (Eickemeyer, Tuttlingen, Germany). The pulse oximeter SurgiVet V90041 (Smiths Medical PM, Inc., Norwell, Massachusetts, USA) continuously records the oxygen saturation, heart rate and photoplethysmogram at the distal M. gastrocnemius of the pigeon. The temperature was kept constant with a water perfused heating blanket and continuously measured rectally with a digital thermometer (Voltcraft Multi-Thermometer DT-300, Conrad Electronic AG, Wollerau, Switzerland).

For the skull thinning surgery we applied 0.5mg/kg of bodyweight Buprenorphine as a preemptive analgesic 30 minutes before the narcosis. The maintenance isoflurane concentration was 2.5 – 3%. HDfUS measurements were performed on the fourth day after surgery at the earliest. To reduce the maintenance isoflurane dose to 1.5 – 2%, 5mg/kg of bodyweight Midazolam was injected 30 minutes before the narcosis in the consequent experiments.

### Trepanation

The bone structure of birds is not solid, but contains many cavities. At each bone/air interface, ultrasound is strongly scattered and reflected because of the high mismatch in the acoustic impedances. Therefore, partial trepanation is inevitable for Power Doppler imaging of the pigeon brain.

After a 25mm long skin incision from rostral to caudal, the skull was cleaned and an area of about (14 mm)^2^ was thinned with a micro drill (K9 EWL type 915, KaVo Dental GmbH, Biberach, Germany). The few hundred μm thick bottom layer of the skull was left intact such that the skull could regrow and the pigeons could fully recover after the experiment. The surgery lasted approximately 2hrs.

### Fixation of the pigeon’s body and head

Wings and legs were mildly attached to the body by appropriately cut sections of hold-up sockings and tape. After that, once positioned horizontally on a body fitting Styrofoam mattress, pigeons were quite calm and did not show any visible signs of stress such as motions or accelerated heartbeat or breathing.

The thinned area of the head was covered with a 3D printed polyamide basin, which was fixed to the skull with dental cement (Harvard Cement normal setting, Harvard Dental International GmbH, Hoppegarten, Germany). To restrict motion to few micrometers, the pigeon’s head was fixated in a custom made stereotactic device with two holders that were designed alongside the basin. The holders were attached to the stereotactic device in place of the common ear plugs and fixated the basin with a key and slot joint. Detailed images of the basin and a 3D model of the holders and basin can be found in the supplementary material (see also Supplementary Fig. 5)

### Orientation in the Pigeon Brain

To guarantee reproducibility of the measurements over days and in view of future comparative pigeon studies we defined the pineal gland in the brain to be the zero location (PG + 0 mm), which is located exactly between the cerebellum and the cerebrum. This defines a well recognizable landmark and is less prone to errors than the usually called upon pigeon stereotactic atlas (Karten, 1967). The transducer angle in all the measurements translates to 65° from the horizontal skull axis in common coordinates.

### Stimulation Sequences

Acoustic stimulation: Two speakers (Stereo Speaker E300, Hama, Monheim, Germany) were placed 0.5 m frontal of the pigeon head. The stimulation sequence consisted of 15 s of hawk or noise sound, followed by 30 s of silence. This sequence was chosen to allow the monitored response to reach its maximum after stimulation started and to fully decay to the baseline after stimulation terminated. The power spectrum of the noise was the same as the hawk sound (cf. supplement material).

Static visual stimulation: Two white LEDs (LL1502HCWW1-CO1 White LED, 6500K by Ledman, Shenzhen, China) were placed approximately 5.5 cm away from the closed eyes of the pigeon in the lateral direction. The stimulation sequence consisted of 15 s on, 30 s off with a flashing frequency of 5 Hz (100 ms on 100 ms off) during the on sequence.

Dynamic visual stimulation: Two 7” TFT monitors (Kkmoon 7” TFT ColorDisplay LED HD, Shenzhen, China) were placed at a distance of 9cm in front of the pigeons head, one at +45° and one at −45° angle to the pigeon axis, thus covering the pigeon’s 90° lateral to 0° frontal field of view (see Supplementary Fig. 6). The stimulation sequence consisted of a white 5 × 5 mm square on a black background, moving within 15 s from the right lateral position to the left where it remained static for 30 s and then moved back to the right. The contrast of the monitors is 500:1.

### Imaging the Brain

For the fUS image acquisition we used a Verasonics Vantage 128 channel system connected to a L22-14v transducer (Verasonics, Kirkland, WA, USA), transmitting at 16MHz and positioned right above the trepanation area. The HDfUS acquisitions were performed multiple times within 4 weeks after surgery. While we observed a small increase in signal attenuation due to back growing bone during this time we still got good signals within the last week without any further drilling. To couple the ultrasound to the skull we used Aquasonic ultrasound transmission gel. The acquisition protocol was executed and processed live in MATLAB 2015a and MATLAB 2016a (MathWorks, Natick, MA, USA).

### Ultrasound sequences

The ultrasound acquisition consisted of angled plane wave imaging (Bercoff et al., 2011; Montaldo et al., 2009) and coherent compounding. Due to ongoing development of the HDfUS algorithm, we used two protocols, using 20 and 36 beams angled between −8° and +8° equidistantly. The ensemble lengths of the two protocols were 300 and 160 continuous with Doppler pulse repetition frequencies of 1 kHz and 300Hz. The maximum actual firing rate achieved was 20 kHz continuously resulting in a data throughput of ∼ 3.3 GB/s, which was processed real time, including image reconstruction and Power Doppler image calculation employing a singular value decomposition filter (Demené et al., 2015). The first (faster) HDfUS protocol was used for the data shown in Fig.1,2,5 and the second protocol for Fig.3,4.

**Fig.1:**
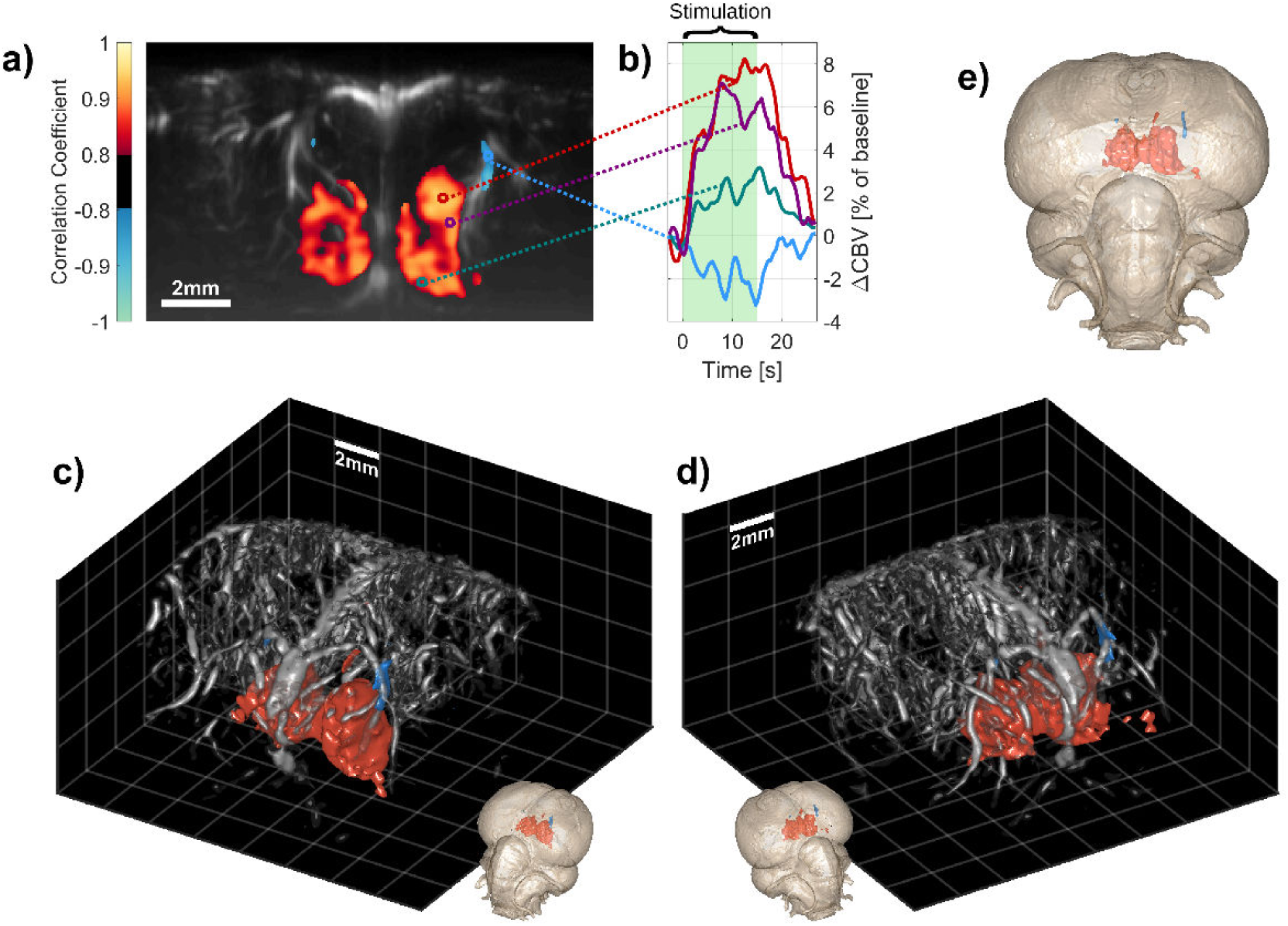
2D and 3D fUS responses in the auditory system. **a)** The Power Doppler image (grey) shows the vascularity of a slice through the pigeon brain. The colored overlay in the image represents the activated regions in the brain evaluated by the Pearson correlation coefficient (cf. methods) over a single stimulus (threshold at |r| > 0.75, red/yellow and blue/green correspond to positive and negative correlation, i.e. CBV increase/decrease). The temporal CBV evolution of four regions of interest (ROIs) of size (200μm)^2^ are shown in **b)** for a single response with the green bar indicating the time of auditory stimulation. **c)** and **d)** 3D reconstructions of the brain activity from a scan of 18 HDfUS acquisitions, separated by 500 μm. The detailed 3D vascularity map (grey) was retrieved by Power Doppler imaging (cf. methods). At the bottom of the 3D images and in **e)**, the activated regions are mapped to an MRI dataset of a pigeon brain. The activity is located in the Field L2 and neighboring areas. The negative CBV change (blue) is located in an arteriole supplying the deeper capillary structure of the brain, where the active regions (i.e. positive CBV change) are situated. See Supplementary Video 1 for the detailed 4D representation of the response.

### Power Doppler processing

In order to filter out the blood component from the backscattered echoes, we decomposed the signal of the coherently compounded images into its singular values over time (Demené et al., 2015) and cut away the tissue and noise signal. The tissue components are represented by the first several singular value decomposition (SVD) modes due to the high backscattering amplitude and static phase, whereas the noise is found in the last modes due to the low intensity. We visually categorized the components relevant for the blood flow to be between 40% and 82% of the SVD modes (i.e. ensemble length), which for instance converts to contributing SVD modes between 120 and 246 for an ensemble length of 300.

### High Definition fUS processing

In HDfUS we were able to continuously process data in real time at up to 20 kHz by using C++/CUDA computing as well as dedicated MATLAB algorithms. Before the filtering, the raw plane wave data was IQ-demodulated, delayed and then beamformed and compounded in the Fourier domain (Kruizinga et al., 2012). The processing was mainly performed on one GPU of a Nvidia Tesla K80 (NVIDIA Corporation, Santa Clara, California, USA). The compounded beamformed frames were continuously saved to a SSD at a framerate of 1 kHz and filtered in parallel to produce the Power Doppler images and evaluate the Pearson correlation coefficients, which facilitates a live evaluation of the acquired data.

### Pearson Correlation Maps

The Pearson correlation **r** is defined to be

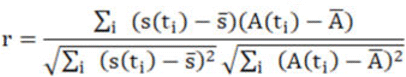

and gives the correlation between the measured blood signal s(t_i_) and the task pattern A(t_i_) at sampling times t_i_. The bar above the coefficients denotes the mean signal. The task pattern A(t_i_) is adapted to the shape of the hemodynamic response (see Fig. 1b). We approximated the response with a trapezoidal shape with a rise time t_rise_ after the stimulus start, a high time t_high_ and a fall time t_fall_. The values for the 15 s hawk stimulus for instance were set to [t_rise_, t_high_, t_fall_] = [7,8,7] s and for the static visual to [t_rise_, t_high_, t_fall_] - [7.5,10,7.5] s.

### 3D Vascularity

The detailed 3D vascularity shown in Fig. 1c/d was retrieved by a separate Power Doppler scan composed of 1026 Power Doppler images at different transducer orientations, following the protocol described in (Demené et al., 2016). The idea of the method is to suppress the point spread function in the elevation direction of the plane wave beam. In order to achieve this, 18 different volumes of the vascularity are acquired by Power Doppler imaging with a rotation of the transducer in z-direction ranging from 0° to 170° in 10° steps. Each volume itself consists of 57 images equidistantly separated by 250 μm in the elevation dimension. By mapping the volumes to a global grid and deconvolving the data, an isotropic resolution of ∼100 μm is achieved in every pixel, resulting in a detailed vascularity map. For the 3D scanning we used a combination of two linear stages (M-UMR8.25, Newport Corporation, CA, USA) connected to a motion controller (MM3000, Newport Corporation, CA, USA) and a rotation stage (M.060.DG, PhysikInstrumente GmbH &Co. KG, Karlsruhe, Germany) connected to a separate motion controller (Mercury C- 863 DC, PhysikInstrumente GmbH &Co. KG, Karlsruhe, Germany). Both controllers are controlled with MATLAB 2016b, so that a full 3D scan may be performed entirely automatically. For the data shown in Supplementary Video 2 and 3, we did not acquire a detailed 3D vascularity map as described above. Here the vascularity was segmented from the Power Doppler image slices making use of 3D skeletonizing (Kerschnitzki et al., 2013) at different thresholds of Power Doppler amplitudes.

### 3D HDfUS Imaging

To reconstruct the 3D activity in the pigeon brain, we acquired several image slices successively. To prevent artefactual signal influences due to long term changes, we first recorded a volume of HDfUS slices in 1mm steps and then recorded another volume, also in 1mm steps but shifted by 0.5mm with respect to the first volume, thus leading to a final spatial out-of-plane resolution of 0.5mm for the combined volume.

For the 3D figures, the data has been spatially filtered with a (150μm)^3^ gaussian smoothing kernel and temporally filtered with a first order Savitzky-Golay of 1.5s (5s for except Fig. 4 to exclude the strong local influence of the cardiac dynamics).

### Reference MRI of the Pigeon Brain

In order to map the fUS dataset to the brain architecture we reference our data to a MRI brain measurement of another pigeon from our loft. With the PG+0 mm point described in the Orientation in the Pigeon Brain section allowed overlaying the datasets. The imaging was conducted with a Biospec 94/20 (Bruker BioSpin MRI GmbH, Germany), a 9.4 T small animal MRI system, at the Institute for Clinical Radiology, OCC Münster, Germany.

## Results

### 4D dynamics in the auditory system

The study of speech acquisition in human infants has strong parallels to the learning process of the song characteristics in some birds (Bolhuis and Gahr, 2006; Kuhl, 2004). In consequence, research on the neural substrates of bird song learning is a prominent model system to examine learning and memory (Bolhuis et al., 2000) and a sensitive and fast neuroimaging of the bird’s auditory system is of great importance. In typical *in vivo* studies, the auditory system of the bird is investigated by electrophysiological recordings (Mandelblat-Cerf et al., 2014) or fMRI (Van Ruijssevelt et al., 2013), which offers very limited field of view and/or spatiotemporal resolution. As a first example of the capabilities of HDfUS we focus therefore on the neural activity evoked by an auditory stimulation in three pigeons.

Figure 1 shows a HDfUS acquisition of one *awake* pigeon, stimulated once for 15s with the sound of an adult hawk scream. Due to the linear architecture of the ultrasound probe, each HDfUS acquisition records the CBV of a slice through the brain continuously (Fig. 1a). The CBV change (Fig. 1b) reveals the shape of the hemodynamic responses within four regions of interest (ROI) evoked by the 15s stimulation. The spatial integration of activation is evaluated by correlating the CBV in each pixel with the task pattern (colored overlay in Fig. 1a). By scanning through the brain in steps of 500 μm with HDfUS acquisitions, the whole three-dimensional response can be appreciated at great detail (Fig. 1c and d). The mapping of the HDfUS datasets to a MRI pigeon brain shows that the activated areas are located within and around the Field L2 (Güntürkün et al., 2013), which is expected due to the associative processing of the known sound pattern. The different ROIs in Fig. 1b furthermore show non-uniform hemodynamic response within the activated area, with signal peaks at different times during the stimulation sequence. There are several possible reasons for the differences in peak time and height between the red, purple and green ROI. First, we expect regions in the middle of the activated area (red and purple) to show a higher peak amplitude than those at the very outer border (green). Second, to the best of our knowledge, this dataset also shows for the first time a drainage of CBV within a supplying larger arteriole during activation (see blue negative values in Fig. 1) and demonstrates the importance of a high spatial resolution in functional neuroimaging. For methods such as fMRI that sample a larger voxel size this drainage effect could potentially lead to a decreased functional signal as the negative and positive change would partly cancel out. Thus, activated brain regions with an associated nearby decrease in CBV are harder to investigate or even identify. For this reason, we would expect the signal peak earlier in regions close to the drained arteriole (red and purple) and later in regions further away (green). Additionally, different functions of these regions could contribute to the differences in hemodynamic response as well.

Our pigeons are naturally accustomed to hawk screaming as they live in an outdoor loft close to a forest and are used both to local and long distance (50-100 km) free flights. In order to evaluate the difference in neural activation between an unknown sound stimulus (noise) and a known sound (hawk), we exposed the awake pigeon to a noise sound of the same frequency spectrum and amplitude as the hawk sound (Fig. 2, sound files are provided in Supplementary Audio). For both cases the response starts similarly within the Field L2, but levels off earlier for the noise case and shows a second feature after the noise sound stops at 15s. This would be expected because there is no variation and/or information during the noise sound, leading to habituation. The only variation causing brain activation is the start and the stop of the noise, thus leading to the two associated peaks approximately five seconds after the start and stop.

**Fig.2:**
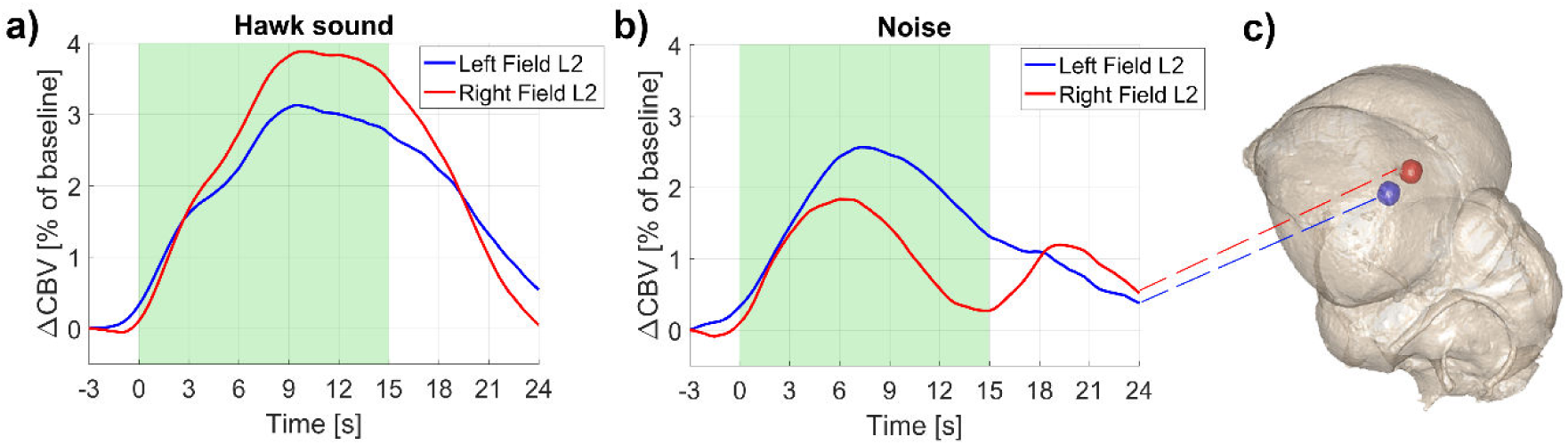
Hawk sound vs. noise in the Field L2. The figure shows the average response in the Field L2 (voxel size 1mm^3^) evoked by a sound carrying information as well as variation (**a**, hawk sound) compared to a noise sound with the same power spectrum (**b,** noise). The data is based on single stimulation sequences (i.e. no error bars). In **c)** the location of the Field L2 is shown in the MRI brain for orientation (ROI position: anterior: PG+5mm, lateral: ±3mm, depth: 5.9mm, size: (1.5mm)^3^).

The activity evoked by the hawk sound was reproduced in all three pigeons (one anaesthetized and two awake) resulting in the same characteristics of the hemodynamic response in terms of shape and localization. However, the amplitude of the signal change in the anaesthetized pigeon was about 50% smaller than in the awake subjects.

### 4D dynamics in the visual system

Along with the auditory system of birds, the neural activity evoked by a visual response is of major importance due to the asymmetric pathway architecture in the pigeon brain (Freund et al., 2016). To investigate the visual system of pigeons, we first recorded HDfUS data of *anaesthetized* pigeons with a non-moving unilaterally flashing light source located in the rear field of view. In Fig. 3 we show the response to a unilateral left eye stimulation of one anaesthetized pigeon with a static light source flashing at 5Hz. The activation is mainly concentrated in the visual wulst of the contralateral hemisphere with a signal increase of up to 5%. The inverted case of right eye stimulation of the same animal also results in an inverted response, but with a 20% higher amplitude (see Supplementary Video 2 and 3).

**Fig.3:**
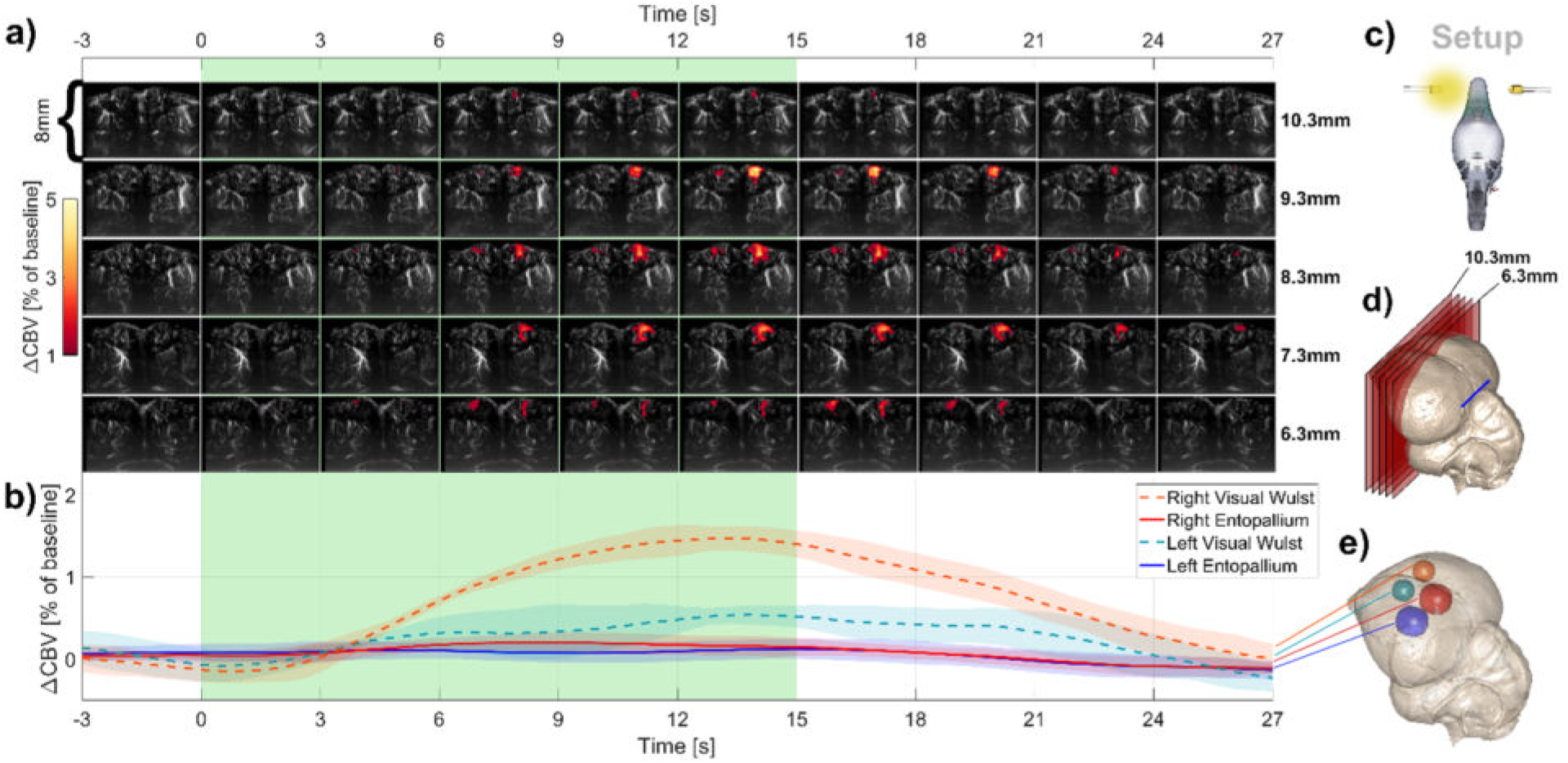
Unilateral left eye visual stimulation of anaesthetized pigeons. **a)** The temporal course of cerebral blood volume change (ΔCBV) within the visual wulst (average over 4 stimulations of one animal) in five brain slices. Each image shows the vasculature as a greyscale map derived from the Power Doppler image and the ΔCBV between 1% and 5% as a colored overlay. The positions of the slices within the brain are shown to the right of the images and in **d)** with respect to the pineal gland (PG) location (blue bar). **b)** The average signal (error band one standard deviation) within four different regions of interest, whose positions are indicated in **e)** with the corresponding colors (ROI position of entopallium: anterior: PG+6.75mm, lateral: ±5.6mm, depth: 4.9mm, size: (3.5mm)^3^; of visual wulst: anterior: PG+8mm, lateral: ±4.6mm, depth: 1.5mm, size: (2.5mm)^3^). The green bar specifies the timespan of stimulation. **c)** A schematic of the setup. See Supplementary Video 2 and 3 for the detailed 4D representation of the response.

With the enhanced temporal resolution offered by HDfUS, the complex dynamics within the activated region can now be studied in greater detail. Fig.4 shows that the area of the highest response is shifted to the posterior of the contralateral visual wulst under right eye stimulation and anterior for the left. The time required to reach the signal peak **s̄_max_** shows that small signal changes tend to correspond to late responses, whereas high s̄_max_ are mainly found around 14s. The responses to visual stimuli in the anaesthetized pigeons were however strongly subjected to individual differences which we trace back to two reasons. First, the strong influence of anesthetics on the neurovascular coupling (Sharp et al., 2015) together with the high maintenance isoflurane concentrations (∼2%) necessary in pigeons (Gunkel and Lafortune, 2005) and second the difficulty to control the eye focus as well as the degree of eyelid closure.

**Fig.4:**
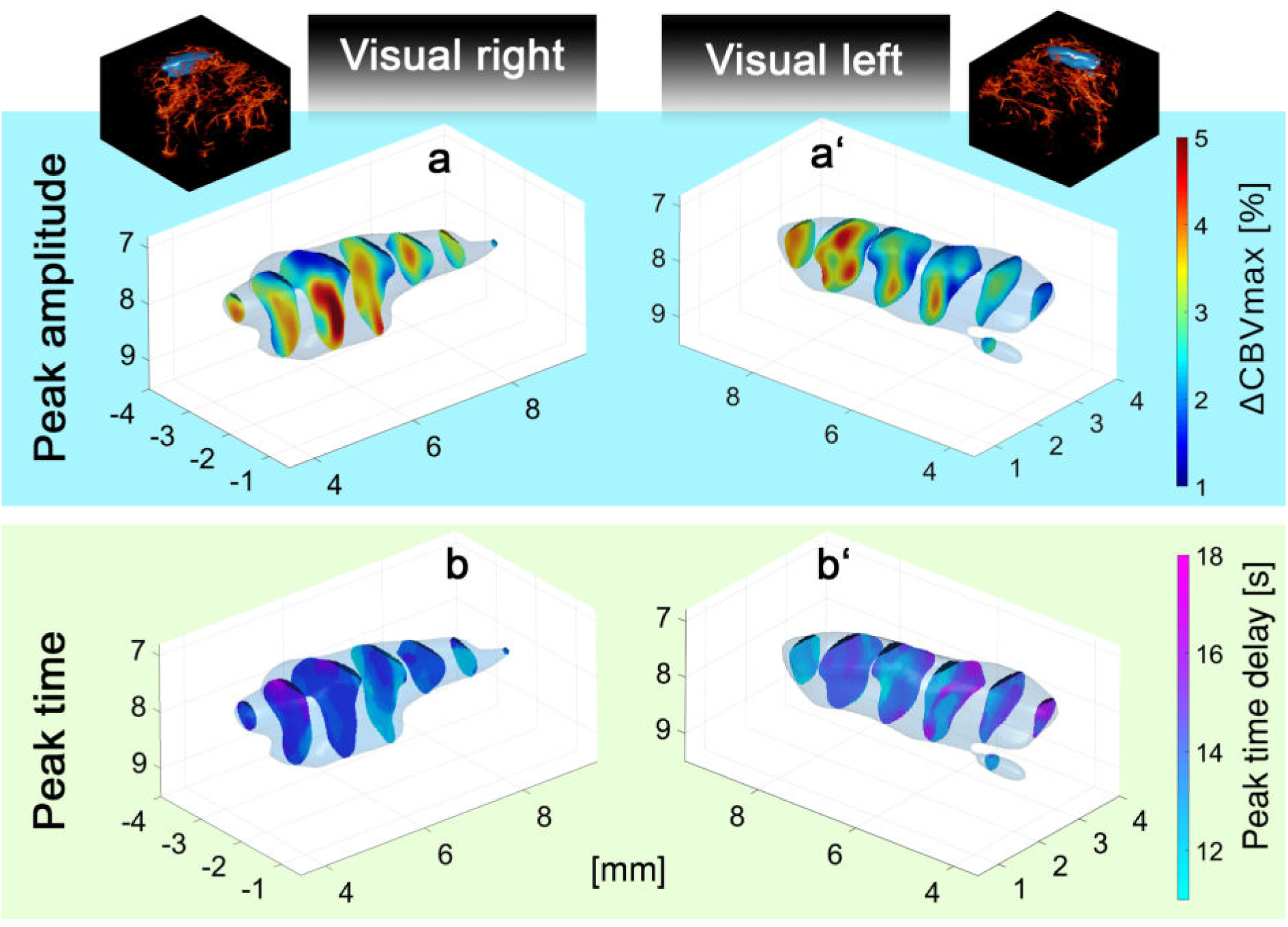
Complex dynamics in the activated regions. The figure shows how the CBV response is spreading in the activated region with the peak amplitude values s̄_max_ in **a/a**’ and the delay time when the peak amplitude s̄_max_ is reached in **b/b**’. The blue activity maps within the vasculature (from Power Doppler imaging) are depicted at the top of the figure for orientation. For a better comprehension of the spatiotemporal spreading in the brain see Supplementary Video 2 and 3.

In order to show the full potential HDfUS provides for neuroscientific questions, we did investigated the visual system in more detail. We performed measurements on two *awake* pigeons stimulated with a moving and constantly illuminating light source. In Fig. 5a-e the neural response within the visual system is shown for a small light horizontally moving from the right to the left in 15s. The resulting response shows a strong dominance of the left entopallium, which is processing the information input from both eyes. The right entopallium is almost solely responding to the contralateral stimulation, which happens at about 7.5s when the dot on the screen starts to appear in the left eye’s field of view. The left hemispheric dominance is also observed with the inverted stimulus pattern (dot on screen moving from left to right) as shown in Fig. 5f. This asymmetry of the pigeon’s visual system has been described before (Freund et al., 2016; Güntürkün et al., 2000), but it has never been shown how the neural activity propagates through different relevant brain areas (i.e. visual wulsts and entopallia) when the pigeon is exposed to a single complex stimulation pattern that excites a wide range of its visual field.

**Fig.5:**
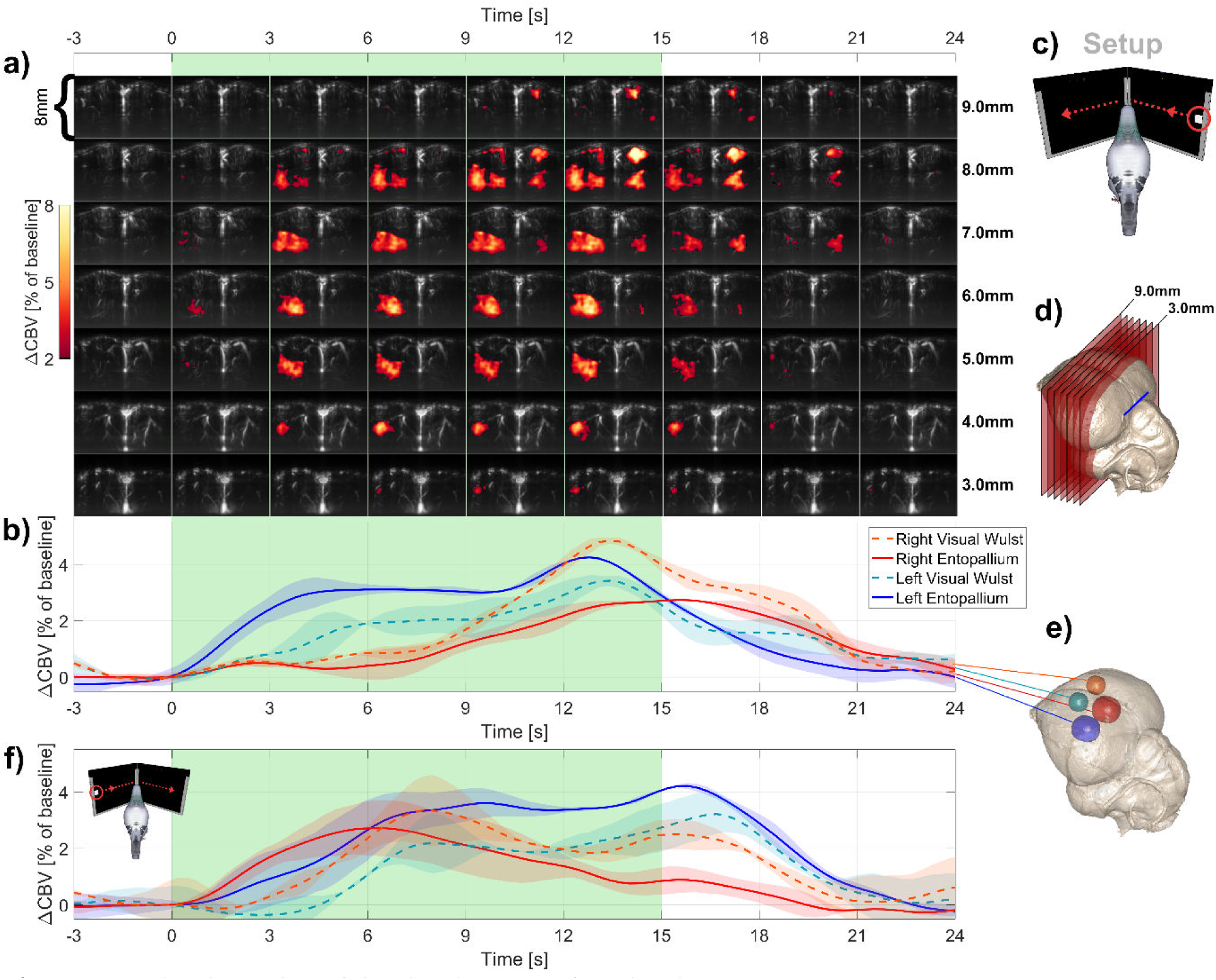
Dynamic stimulation of the visual system of awake pigeons. **a)** The temporal course of ΔCBV within the visual wulst and entopallium (average over two stimulations of one animal) in seven brain slices. Each image shows the vasculature as a greyscale map derived from the Power Doppler image and the ΔCBV between 2% and 8% as a colored overlay. The positions of the slices within the brain are shown to the right of the images and in **d)** with respect to the pineal gland (PG) location (blue bar). **b)** The average signal within four different regions of interest, whose positions are indicated in **e)** with the corresponding colors. The green bar specifies the timespan of stimulation. **c)** Schematic of the setup. **f)** The response to the inverted stimulation pattern with the light source moving from left to right. The left hemispheric dominance is apparent for the two cases. See Supplementary Video 4 and 5 for the detailed 4D representation of the response.

## Discussion and conclusion

In this study, we introduce a new data acquisition scheme which pushes current fUS technology to enhanced spatiotemporal resolution, enabling for the first time successful studies in species such as pigeons with unfavorably low heartbeat. With the resolution reached by HDfUS, we determined the activated regions within the pigeon brain evoked by various stimulations and setups relevant in the field of avian neurobiology. Furthermore, exploiting the high spatiotemporal resolution offered by HDfUS, we were able to resolve locally separated temporal spreading of the hemodynamic response due to a complex stimulation such as a moving light source. This was not revealed previously by any other method. We believe that he implications and potential of this method for neuroimaging studies in avian species are huge. It could for example lead to a better understanding of interhemispheric communication and the neural processing associated with language learning and vocalization, which is of relevance for developmental disorders and language tasks in humans.

While these are the first functional ultrasound measurements of a non-mammalian species as well as the first ones on an animal without a cortical brain architecture, the enhanced fUS sensitivity offered by HDfUS may also lead to a better understanding of the hemodynamic responses in the rodent brain. Especially, HDfUS paves the way for studying weaker or faster responses when an effective suppression of the heartbeat influence at an enhanced spatiotemporal resolution is crucial. With respect to fUS of the human brain, an effective suppression of the heartbeat influence is even more crucial due to the stronger fluctuation of the CBV compared to small animals.

Furthermore, the detailed revelation of the 3D dynamics demonstrates great potential in understanding the hemodynamic response more thoroughly. Even though the HDfUS measurement is still very time consuming due to the use of 1D transducer arrays that require subsequent scanning of single 2D brain slices in order to obtain 3D spatial information, the development of 2D transducer arrays will provide the necessary step forward. Two dimensional arrays already show excellent results of directly recording 4D dynamics in larger vessels (Provost et al., 2014). An expansion of the technology to small animal imaging in the near future is foreseeable. Along with the fUS studies on awake and mobile rodents (Sieu et al., 2015; Urban et al., 2015), this shows the great potential of fUS imaging to become a new standard method revealing unknown hemodynamics at excellent SNR and spatiotemporal resolution. In addition, unlike fMRI, fUS is essentially a portable method which leaves almost unlimited accessible experimental space around the animal allowing for unconstrained design of the animal’s environment and panoramic perception pattern.

## Acknowledgements

We would like to thank C.Faber and co-workers of the University of Münster, Germany, for the MRI data of pigeon brain.

## References

Bercoff, J., Montaldo, G., Loupas, T., Savery, D., Meziere, F., Fink, M., Tanter, M., 2011. Ultrafast compound doppler imaging: providing full blood flow characterization. IEEE Trans. Ultrason. Ferroelectr. Freq. Control 58, 134–147. https://doi.org/10.1109/TUFFC.2011.1780

Bolhuis, J.J., Gahr, M., 2006. Neural mechanisms of birdsong memory. Nat. Rev. Neurosci. 7, 347–357. https://doi.org/10.1038/nrn1904

Bolhuis, J.J., Zijlstra, G.G.O., den Boer-Visser, A.M., der Zee, E.A.V., 2000. Localized neuronal activation in the zebra finch brain is related to the strength of song learning. Proc. Natl. Acad. Sci. 97, 2282–2285. https://doi.org/10.1073/pnas.030539097

Browning, R., Bruce Overmier, J., Colombo, M., 2011. Delay activity in avian prefrontal cortex – sample code or reward code? Eur. J. Neurosci. 33, 726–735. https://doi.org/10.1111/j.1460-9568.2010.07540.x

Demene, C., Baranger, J., Bernal, M., Delanoe, C., Auvin, S., Biran, V., Alison, M., Mairesse, J., Harribaud, E., Pernot, M., Tanter, M., Baud, O., 2017. Functional ultrasound imaging of brain activity in human newborns. Sci. Transl. Med. 9, eaah6756. https://doi.org/10.1126/scitranslmed.aah6756

Demené, C., Deffieux, T., Pernot, M., Osmanski, B.-F., Biran, V., Gennisson, J.-L., Sieu, L.-A., Bergel, A., Franqui, S., Correas, J.-M., Cohen, I., Baud, O., Tanter, M., 2015. Spatiotemporal Clutter Filtering of Ultrafast Ultrasound Data Highly Increases Doppler and fUltrasound Sensitivity. IEEE Trans. Med. Imaging 34, 2271–2285. https://doi.org/10.1109/TMI.2015.2428634

Demené, C., Tiran, E., Sieu, L.-A., Bergel, A., Gennisson, J.L., Pernot, M., Deffieux, T., Cohen, I., Tanter, M., 2016. 4D microvascular imaging based on ultrafast Doppler tomography. NeuroImage 127, 472–483. https://doi.org/10.1016/j.neuroimage.2015.11.014

Doupe, A.J., Kuhl, P.K., 1999. Birdsong and human speech: Common Themes and Mechanisms. Annu. Rev. Neurosci. 22, 567–631. https://doi.org/10.1146/annurev.neuro.22.1.567

Freund, N., Valencia-Alfonso, C.E., Kirsch, J., Brodmann, K., Manns, M., Güntürkün, O., 2016. Asymmetric top-down modulation of ascending visual pathways in pigeons. Neuropsychologia 83, 37–47. https://doi.org/10.1016/j.neuropsychologia.2015.08.014

Gesnik, M., Blaize, K., Deffieux, T., Gennisson, J.-L., Sahel, J.-A., Fink, M., Picaud, S., Tanter, M., 2017. 3D Functional Ultrasound Imaging of the cerebral visual system in rodents. NeuroImage. https://doi.org/10.1016/j.neuroimage.2017.01.071

Gunkel, C., Lafortune, M., 2005. Current Techniques in Avian Anesthesia. Semin. Avian Exot. Pet Med., Anesthesia and Analgesia 14, 263–276. https://doi.org/10.1053/j.saep.2005.09.006

Güntürkün, O., Diekamp, B., Manns, M., Nottelmann, F., Prior, H., Schwarz, A., Skiba, M., 2000. Asymmetry pays: visual lateralization improves discrimination success in pigeons. Curr. Biol. 10, 1079–1081. https://doi.org/10.1016/S0960-9822(00)00671-0

Güntürkün, O., Verhoye, M., Groof, G.D., der Linden, A.V., 2013. A 3-dimensional digital atlas of the ascending sensory and the descending motor systems in the pigeon brain. Brain Struct. Funct. 218, 269–281. https://doi.org/10.1007/s00429-012-0400-y

Hedrich, H.J., 2012. The Laboratory Mouse. Academic Press.

Jarvis, E.D., 2004. Learned Birdsong and the Neurobiology of Human Language. Ann. N. Y. Acad. Sci. 1016, 749–777. https://doi.org/10.1196/annals.1298.038

Jensen, J.A., 1996. Estimation of Blood Velocities Using Ultrasound: A Signal Processing Approach. Cambridge University Press.

Jung-Beeman, M., 2005. Bilateral brain processes for comprehending natural language. Trends Cogn. Sci. 9, 512–518. https://doi.org/10.1016/j.tics.2005.09.009

Karten, P.H.J., 1967. A Stereotaxic Atlas of the Brain of the Pigeon. The Johns Hopkins University Press.

Kerschnitzki, M., Kollmannsberger, P., Burghammer, M., Duda, G.N., Weinkamer, R., Wagermaier, W., Fratzl, P., 2013. Architecture of the osteocyte network correlates with bone material quality. J. Bone Miner. Res. 28, 1837–1845. https://doi.org/10.1002/jbmr.1927

Kruizinga, P., Mastik, F., Jong, N. de, van der Steen, A.F.W., Soest, G. van, 2012. Plane-wave ultrasound beamforming using a nonuniform fast fourier transform. IEEE Trans. Ultrason. Ferroelectr. Freq. Control 59. https://doi.org/10.1109/TUFFC.2012.2509

Kuhl, P.K., 2004. Early language acquisition: cracking the speech code. Nat. Rev. Neurosci. 5, 831–843. https://doi.org/10.1038/nrn1533

Logothetis, N.K., 2008. What we can do and what we cannot do with fMRI. Nature 453, 869–878. https://doi.org/10.1038/nature06976

Macé, E., Montaldo, G., Cohen, I., Baulac, M., Fink, M., Tanter, M., 2011. Functional ultrasound imaging of the brain. Nat. Methods 8, 662–664. https://doi.org/10.1038/nmeth.1641

Macé, E., Montaldo, G., Osmanski, B.F., Cohen, I., Fink, M., Tanter, M., 2013. Functional ultrasound imaging of the brain: theory and basic principles. IEEE Trans. Ultrason. Ferroelectr. Freq. Control 60, 492–506. https://doi.org/10.1109/TUFFC.2013.2592

Mandelblat-Cerf, Y., Las, L., Denisenko, N., Fee, M.S., 2014. A role for descending auditory cortical projections in songbird vocal learning. eLife 3. https://doi.org/10.7554/eLife.02152

Manns, M., Römling, J., 2012. The impact of asymmetrical light input on cerebral hemispheric specialization and interhemispheric cooperation. Nat. Commun. 3, 696. https://doi.org/10.1038/ncomms1699

Montaldo, G., Tanter, M., Bercoff, J., Benech, N., Fink, M., 2009. Coherent plane-wave compounding for very high frame rate ultrasonography and transient elastography. IEEE Trans. Ultrason. Ferroelectr. Freq. Control 56, 489–506. https://doi.org/10.1109/TUFFC.2009.1067

Ng, B.S.W., Grabska-Barwińska, A., Güntürkün, O., Jancke, D., 2010. Dominant vertical orientation processing without clustered maps: early visual brain dynamics imaged with voltage-sensitive dye in the pigeon visual Wulst. J. Neurosci. Off. J. Soc. Neurosci. 30, 6713–6725. https://doi.org/10.1523/JNEUROSCI.4078-09.2010

Niranjan, A., Christie, I.N., Solomon, S.G., Wells, J.A., Lythgoe, M.F., 2016. fMRI mapping of the visual system in the mouse brain with interleaved snapshot GE-EPI. NeuroImage 139, 337–345. https://doi.org/10.1016/j.neuroimage.2016.06.015

Osmanski, B.-F., Pezet, S., Ricobaraza, A., Lenkei, Z., Tanter, M., 2014. Functional ultrasound imaging of intrinsic connectivity in the living rat brain with high spatiotemporal resolution. Nat. Commun. 5, 5023. https://doi.org/10.1038/ncomms6023

Provost, J., Papadacci, C., Arango, J.E., Imbault, M., Fink, M., Gennisson, J.-L., Tanter, M., Pernot, M., 2014. 3D ultrafast ultrasound imaging in vivo. Phys. Med. Biol. 59, L1–L13. https://doi.org/10.1088/0031-9155/59/19/L1

Rungta, R.L., Osmanski, B.-F., Boido, D., Tanter, M., Charpak, S., 2017. Light controls cerebral blood flow in naive animals. Nat. Commun. 8, 14191. https://doi.org/10.1038/ncomms14191

Sharp, P.S., Shaw, K., Boorman, L., Harris, S., Kennerley, A.J., Azzouz, M., Berwick, J., 2015. Comparison of stimulus-evoked cerebral hemodynamics in the awake mouse and under a novel anesthetic regime. Sci. Rep. 5, 12621. https://doi.org/10.1038/srep12621

Shung, K.K., Sigelmann, R.A., Reid, J.M., 1976. Scattering of Ultrasound by Blood. IEEE Trans. Biomed. Eng. BME-23, 460–467. https://doi.org/10.1109/TBME.1976.324604

Sieu, L.-A., Bergel, A., Tiran, E., Deffieux, T., Pernot, M., Gennisson, J.-L., Tanter, M., Cohen, I., 2015. EEG and functional ultrasound imaging in mobile rats. Nat. Methods 12, 831–834. https://doi.org/10.1038/nmeth.3506

Tanter, M., Fink, M., 2014. Ultrafast imaging in biomedical ultrasound. IEEE Trans. Ultrason. Ferroelectr. Freq. Control 61, 102–119. https://doi.org/10.1109/TUFFC.2014.2882

Tiran, E., Ferrier, J., Deffieux, T., Gennisson, J.-L., Pezet, S., Lenkei, Z., Tanter, M., 2017. Transcranial Functional Ultrasound Imaging in Freely Moving Awake Mice and Anesthetized Young Rats without Contrast Agent. Ultrasound Med. Biol. https://doi.org/10.1016/j.ultrasmedbio.2017.03.011

Urban, A., Dussaux, C., Martel, G., Brunner, C., Mace, E., Montaldo, G., 2015. Real-time imaging of brain activity in freely moving rats using functional ultrasound. Nat. Methods 12, 873–878. https://doi.org/10.1038/nmeth.3482

Urban, A., Mace, E., Brunner, C., Heidmann, M., Rossier, J., Montaldo, G., 2014. Chronic assessment of cerebral hemodynamics during rat forepaw electrical stimulation using functional ultrasound imaging. NeuroImage 101, 138–149. https://doi.org/10.1016/j.neuroimage.2014.06.063

Van Ruijssevelt, L., Van der Kant, A., De Groof, G., Van der Linden, A., 2013. Current state-of-the-art of auditory functional MRI (fMRI) on zebra finches: Technique and scientific achievements. J. Physiol.-Paris, Special issue: Physiological mechanisms of song learning and production 107, 156–169. https://doi.org/10.1016/j.jphysparis.2012.08.005

Voss, H.U., Tabelow, K., Polzehl, J., Tchernichovski, O., Maul, K.K., Salgado-Commissariat, D., Ballon, D., Helekar, S.A., 2007. Functional MRI of the zebra finch brain during song stimulation suggests a lateralized response topography. Proc. Natl. Acad. Sci. 104, 10667–10672. https://doi.org/10.1073/pnas.0611515104

